# Improving DNA Modeling with WaveDNA: Enhancing Speed, Generalizability, and Interpretability through Wavelet Transformation

**DOI:** 10.1101/2025.11.07.687194

**Authors:** Lorenzo Ruggeri, Manuel Tognon, Rosalba Giugno

## Abstract

Transcription factors (TFs) regulate gene expression by binding to short, specific DNA sequences, known as transcription factor binding sites (TFBSs). Accurate identification of TFBSs is fundamental for understanding transcriptional regulation. By leveraging their ability to capture complex non-linear patterns and hierarchical dependencies underlying TF-DNA binding deep learning (DL) has emerged as the state-of-the-art approach for modeling and identifying TFBSs. However, current models often require extensive pretraining, involve large parameter sets, and offer limited interpretability. To address these limitations, we introduce WaveDNA, a lightweight and interpretable DL framework that encode DNA sequences into two-dimensional representations using wavelet transforms. This approach enables the use of convolutional neural networks pretrained on images, facilitating efficient transfer learning without requiring large-scale genomic data pretraining. Across diverse ENCODE ChIP-seq datasets spanning different TFs, WaveDNA achieves predictive accuracy comparable to state-of-the-art DL models while using approximately fivefold fewer parameters and substantially less computational resources. Moreover, representing DNA sequences as images allows the direct application of established computer vision interpretability techniques to visualize the learned binding patterns. Together, these results demonstrate that WaveDNA offers a scalable, computationally efficient, and interpretable alternative for modeling TF-DNA interactions.

## Introduction

Transcription factors (TFs) are key regulatory proteins that control the transcriptional states of cells governing their identity and function [1]. TFs exert their regulatory function by binding to short (typically ∼10 nucleotides long [2]), specific DNA sequences, known as transcription factor binding sites (TFBSs). TFBSs are characterized by recurring, similar, but not identical sequence patterns, referred to as motifs. The ability of TFs to regulate gene expression hinges on their ability to recognize and selectively bind motifs. This process is influenced by both sequence nucleotide composition and chromatin context [3]. Consequently, the accurate discovery and modeling of TFBS motifs (motif discovery) is fundamental to unraveling the mechanisms of transcriptional regulation and remains a pivotal challenge in computational biology [4]. Over the years, different experimental methods have been developed to map TFBSs, both in vitro and in vivo. The advent of high-throughput techniques, including protein binding micro-arrays (PBMs) [5], SELEX [6], and ChIP-seq [7], has enabled genome-wide profiling of TF binding. Despite limitations in resolution and indirect binding artifacts, ChIP-seq has become the ‘gold standard’ for in vivo TFBS identification [1]. Complementary methods, such as DNase-seq [8] and ChIP-exo [9] have been proposed to refine binding localization and address some of these limitations. To extract TFBS motifs from experimental datasets, computational modeling has played a key role. Position weight matrices (PWMs) [10] have long been the standard approach due to their simplicity and interpretability, as they represent nucleotide preferences at each motif position. However, their assumption of positional independence and lack of sequence context can limit predictive accuracy, particularly in complex regulatory contexts [4]. To address these limitations, machine learning models, such as support vector machines (SVMs), have employed to identify and model motifs [11]. By leveraging k-mer-based representations, these models overcome the independence assumption and include sequence context information into the predictive framework. Although SVMs often outperform PWMs in classification tasks, they are constrained by fixed k-mer lengths and are highly dependent on the quality of training data [12]. Over the past decade, deep learning (DL) methods have emerged as the state-of-the-art tools for learning and discovering TFBSs [13]. DL models excel at capturing the complex, hierarchical dependencies that govern TF–DNA interactions within genomic sequences. Among them, convolutional neural networks (CNNs) have been widely adopted for motifs discovery and modeling [14, 15, 16, 17]. CNNs apply non-linear transformations to input sequences and learn rich, high-dimensional representations of binding patterns, enabling accurate predictions of TFBSs. Typically, genomic sequences are encoded as one- or two-dimensional arrays with four channels corresponding to nucleotide bases (A, C, G, T), framing TFBS prediction as a binary image classification task [14]. Due to their local connectivity and parameter-sharing design, CNNs are computationally efficient and effective at detecting local sequence patterns like TF motifs. However, their limited receptive field makes it challenging to capture long-range dependencies between distal regulatory elements, which often play a critical role in TF-DNA binding events [18, 19]. Although techniques, such as dilated convolutions or deeper architectures can partially address this by expanding the context window, they often increase model complexity and reduce interpretability [20]. Recently, transformer models [21] have been applied to regulatory genomics. By leveraging self-attention mechanisms, transformers can model global dependencies across entire sequences, generating contextualized representations that capture complex interactions between proximal and distal genomic elements, such as enhancers, promoters, and insulators. Thus, transformers are well-suited for regulatory genomics, where long-range interactions play a crucial role in gene expression control. Different genomic transformer models have emerged, pretrained on large-scale genomic sequence data using self-supervised objectives and fine-tuned for specific downstream tasks. Models such as Enformer [22], GET [23], and Basenji [24] combine convolutional layers with attention mechanisms to predict regulatory signals across long input DNA sequences. In parallel, masked language models (MLMs) such as DNABERT [25, 26] and Nucleotide Transformer (NT) [27] have been specifically designed and fine-tuned for genomic classification tasks. While transformer models have demonstrated remarkable performance in genomic sequence modeling, they are not without limitations. They typically require extensive pretraining on large-scale, domain-specific datasets to learn effective representations, which may be a significant hurdle in genomics, where labeled data are often limited and costly to obtain. Moreover, transformers are characterized by a very large number of parameters, often in the range of hundreds of millions to billions, which increases both training time and the risk of overfitting in low-data regimes [28]. Their self-attention mechanism also scales quadratically with sequence length, making them computationally expensive and memory-intensive [29, 30]. Finally, interpretability remains a key challenge: the complexity of multi-head self-attention across numerous layers makes it difficult to extract biologically meaningful insights from model predictions [31]. Recently, HyenaDNA [32], a genomic foundation model based on implicit convolutions, has been proposed as a scalable alternative to transformers. It achieves sub-quadratic scaling with sequence length and significantly reduces training time while maintaining full global context at each layer, addressing scalability limitations of transformers. Building on this, we introduce WaveDNA, a novel framework that revisits CNNs as a scalable, memory-efficient alternative to transformer-based architectures for genomic sequence modeling and TF binding prediction tasks. By using wavelet transforms to encode DNA sequences into time-frequency representations, WaveDNA models capture both local and global features. This encoding transforms genomic sequences into multi-scale image-like inputs, unlocking the opportunity of using pretrained image-based CNN architectures, such as ResNet [33], via transfer learning. Thus, WaveDNA does not require extensive pretraining on genomic data and enables effectively generalization even with limited labeled data. Moreover, representing DNA sequences as images allows model interpretation by leveraging established feature visualization techniques from computer vision. We compared the performance of WaveDNA and three state- of-the-art DL models, HyenaDNA [32], DNABERT-2 [26], and Nucleotide Transformer (NT) [27], on TFBS prediction tasks using 13 TF ChIP-seq datasets from ENCODE [34]. The selected TFs span a range of DNA-binding domains, motif lengths, information content, and sequence variability, ensuring a comprehensive evaluation represented across distinct binding preferences and regulatory context. Our results demonstrate that WaveDNA achieves predictive accuracy comparable to transformer-based models while using substantially fewer computational resources and without the need for explicit pretraining on genomic data. In addition, WaveDNA enables improved interpretability, as it allows to directly adapt established techniques from computer vision to DNA sequences, such as Grad-CAM [35]. In the following sections, we introduce our novel method WaveDNA, detailing its architecture. We then provide an overview of the three DL models (HyenaDNA, DNABERT-2, and NT) used for benchmarking, highlighting their key features. Subsequently, we describe the datasets employed in our experiments. We also outline the training procedures, hyperparameter configurations, and evaluation metrics used in the study. Then, we detail the systematic evaluation of WaveDNA and the baseline models using multiple accuracy metrics. Beyond predictive accuracy, we examine the models’ computational requirements, such as training time and memory consumption. Furthermore, we examine the interpretability of WaveDNA’s outputs by visualizing learned representations and comparing them with experimentally validated ChIP-seq data. Through this comprehensive evaluation we show that by integrating wavelet-based sequence encoding and transfer learning, CNN architectures can provide a valuable lightweight alternative for modeling TF-DNA binding events while overcoming the scalability and interpretability limitations of transformers.

## Materials and Methods

WaveDNA is a deep learning framework for predicting transcription factor binding sites (TFBS) by transforming raw DNA sequences into image-like representations using Continuous Wavelet Transforms (CWT) [36].The framework comprises three main components: (i) encoding nucleotide sequences into two-dimensional frequency-domain representations with the CWT (Figure 1A),(ii) classifying the resulting wavelet images with a residual CNN (ResNet50) (Figure 1B), and (iii) model’s decision-making process interpretability (Figure 1C). We first detail the CWT-based encoding process and the CNN architecture for classification, followed by the interpretability module based on Gradient-weighted Class Activation Mapping (Grad-CAM) [35]. Then, we describe the datasets used for model training, validation, and benchmarking. Finally, we introduce the state-of-the-art DL methods used to benchmark the performance of WaveDNA across multiple evaluation metrics.

**Fig. 1:**
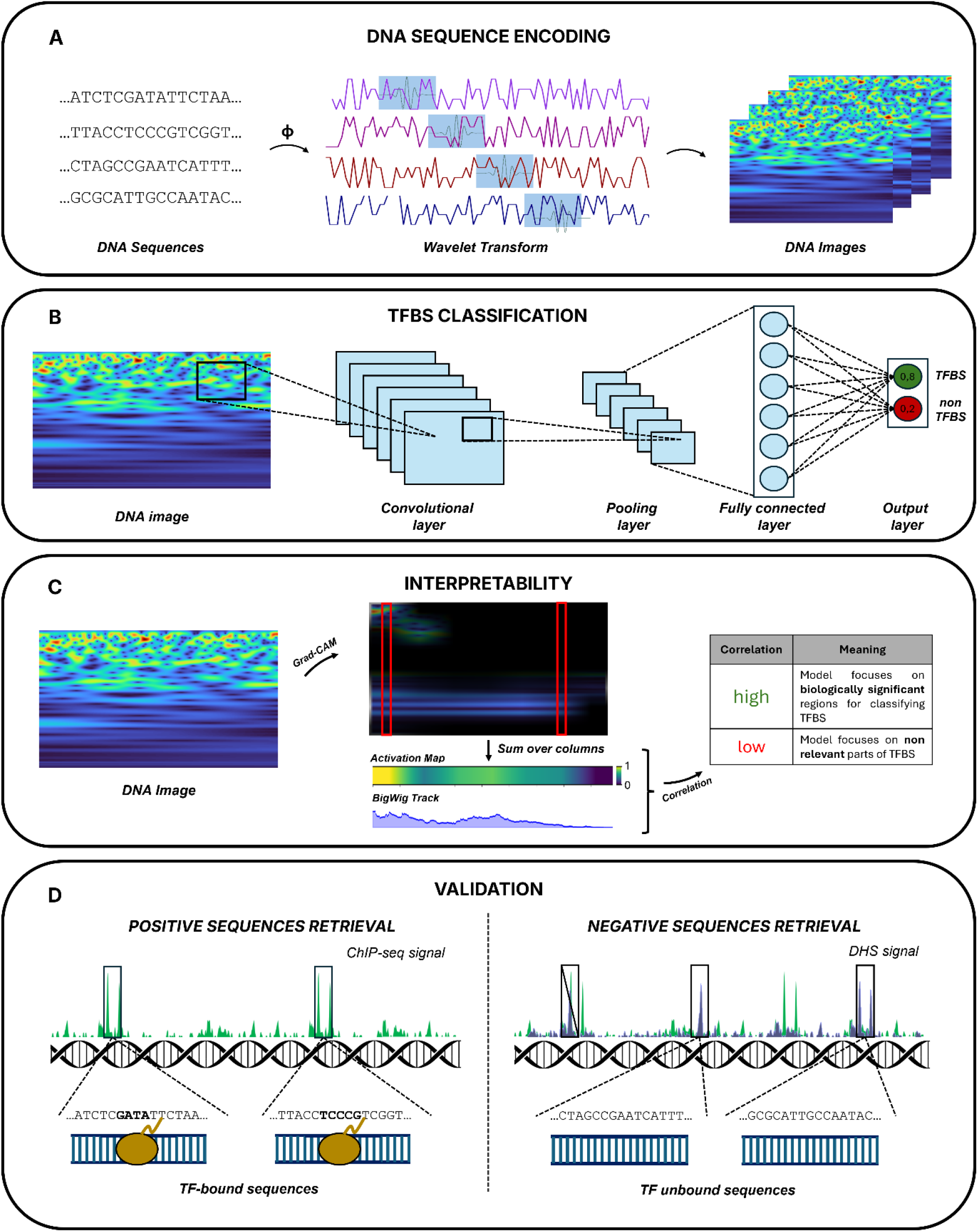
Overview of the WaveDNA framework for TFBSs prediction. WaveDNA integrates signal processing and deep learning to predict TFBSs directly from raw DNA sequences. (A) Each DNA sequence is converted into a one-dimensional discrete signal through a deterministic numerical mapping ϕ: {A, C, G, T} → {1, 2, 3, 4}. The resulting signal is transformed into a two-dimensional representation using the continuous wavelet transform with a Morlet mother wavelet. The coefficient matrices obtained capture signal intensity across both scales and sequence positions. (B) The generated wavelet images are processed by a 50-layer ResNet50 convolutional neural network, which outputs confidence scores classifying each sequence as bound or unbound by the target TF. (C) Model interpretability is assessed using Grad-CAM to identify regions that most influence TFBS predictions. Activation maps are projected onto the sequence axis to produce one-dimensional profiles which are compared with experimental ChIP-seq BigWig tracks. (D) Model validation uses positive samples overlapping ChIP-seq peaks of the target TF (left) and negative sequences drawn from DNase I hypersensitive regions that do not overlap any ChIP-seq peaks (right).

### From Sequence to Image: Sequence Encoding via Continuous Wavelet Transform

In the encoding phase, each DNA sequence is first converted into a one-dimensional numerical signal by applying a scalar mapping function:

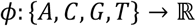

with the nucleotide-to-scalar assignments defined as

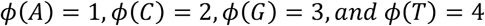

The mapping yields a discrete signal *x*[*n*] that preserves the original nucleotide order, while making it suitable for applying signal processing techniques (**Figure 1A**). To extract both spatial (local) and frequency (global) patterns from *x*[*n*], WaveDNA applies the CWT using the Morlet wavelet as the mother function [36]. Unlike the Fourier transform, which provides a global frequency representation of the entire signal and therefore lacks spatial or temporal specificity, the wavelet transform decomposes the signal into localized basis functions that vary in both scale and position. This allows for the simultaneous analysis of frequency and location, enabling systematic examination of transient signal features, such as peaks or discontinuities across varying scales, while preserving their positional information. Formally, given a signal *S*(*t*), the CWT computes the inner product between it and a mother wavelet *M*(*σ*, Δ), scaled by *σ* and shifted by Δ [37]:

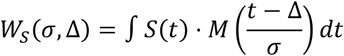

where *σ* and Δ define the wavelet’s dilation and translation, respectively. The resulting wavelet coefficients *W*_*s*_(*σ*, Δ) form a two-dimensional matrix representing the similarity between the signal and the wavelet at different locations and frequencies. These matrices are then rendered as images by mapping coefficient magnitudes to pixel intensities, forming an interpretable and compact visual representation of DNA sequence suitable for CNN input.

### Classification via Residual Convolutional Neural Networks

The generated wavelet images are subsequently processed using ResNet50 [38], a 50-layers deep CNN architecture composed of multiple residual blocks (**Figure 1B**). Each block includes shortcut connections that adds the block’s input to its output, enabling the network to learn identity mappings and mitigating issues related to vanishing gradients. The output of each residual block is computed as:

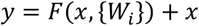

where *x* is the input to the block, *F* denotes a function composed of convolutional, batch normalization, and activation layers, and *W*_*i*_ are the learning weights. This architecture facilitates gradient flow and promotes the reuse of features between layers. For the TFBS prediction task, the final classification layers is a fully connected layer with two output neurons representing the binary classes: TF-bound and unbound sequences.

### Model Interpretability via Grad-CAM

To provide insights into the model’s decision-making process, WaveDNA employs Grad-CAM [35], an extension of the original CAM method [39] (**Figure 1C**). Grad-CAM overcomes limitations of earlier methods by producing class-discriminative, high-resolution localization maps, even in complex architectures with complex learning objectives and fully connected layers. Given an input image and a target class score *c*, Grad-CAM computes the gradients of the class score *y*^*c*^ with respect to the activation maps *A*^*k*^ *∈* ℝ^*u*×*v*^ of a convolutional layer:

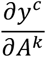

Gradients are globally averaged over the spatial dimensions to obtain the neuron importance weights:

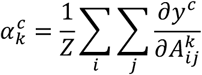

where *Z* = *u* × *v* is the number of spatial positions in the feature map. These weights reflect the contribution of each feature map *A*^*k*^ to the target class. The class-discriminative localization map 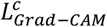is computed as:

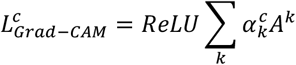

In WaveDNA, Grad-CAM is applied to the convolutional layer exhibiting the highest activation variance for each dataset. The resulting heatmaps highlight regions of the wavelet image, and thus of the original DNA sequence, that contribute most strongly to the model’s classification as TF-bound or unbound. These visualizations not only serve to validate the model’s predictions, but also enable biological interpretation by identifying enriched or recurrent features, such as sequence motifs or periodic patterns, across genomic sequences.

### Validation Datasets

To evaluate the performance of WaveDNA in TFBS prediction, we constructed comprehensive datasets using experimentally validated genomic data from the ENCODE database [34] (**Supplementary File Section 1**). Positive sequence data sets were constructed by retrieving optimal IDR-threshold ChIP sequence peaks for 13 human TFs, spanning different DNA-binding domain classes defined by Lambert et al. [1]. These peaks represent high-confidence binding events, serving as positive samples for training and testing (**Figure 1D**). To construct negative sequence datasets, we used DNase I hypersensitive sites (DHSs) retrieved from different human tissues [40], which mark open chromatin regions accessible to TF binding. To ensure that negative sequences were not bound by the TF of interest, DHS sites were systematically filtered to exclude any site partially or completely overlapping with ChIP-seq peaks. This conservative filtering strategy yielded negative sequences that are chromatin-accessible yet unbound by the target TF, thereby increasing the classification difficulty. To robustly evaluate model generalization and reduce the risk of chromosome-specific overfitting, we implemented a chromosome-stratified cross-validation strategy (**Supplementary File Section 1**).

For each TF, chromosomes 1, 2, 3, 6, and 17 were iteratively held out for testing, while the remaining chromosomes were used for training. These chromosomes were chosen for their high coverage and sequence diversity (**Supplementary Figures 1 and 2**). This chromosome-stratified scheme is widely adopted in genomic benchmarks [41, 42], as it preserves independence between training and test datasets and provides a robust approximation of real-world predictive performance. To address the potential performance inflation caused by homologous sequences between training and test sets [43], we assessed sequence similarity across all datasets. We found that homologous sequences represented < 1% in all experiments, suggesting minimal risk of performance inflation caused by homology (**Supplementary File Section 2 and Supplementary Table 1**). The resulting datasets provide a robust and realistic framework for evaluating WaveDNA and state-of-the-art models on TFBS prediction tasks.

### Benchmark Models for TFBS Prediction

We evaluated the performance of WaveDNA against DL models spanning both convolutional neural network (CNN)- and transformer-based architectures. For each model, we summarize their core architectural features and previous performance in relevant genomic tasks. Supplementary File Section 3 provides detailed descriptions of the compared methods.

#### Convolutional Neural Networks

CNNs are among the earliest DL architectures applied to TFBS prediction. CNNs process one-hot encoded DNA sequences, employing convolutional filters to slide across the input and detect spatially localized patterns, such as sequence motifs [14]. Through these convolutional filters, CNNs extract position-invariant features by detecting recurrent subsequences, producing activation maps that reflect motif occurrences along the sequence. Subsequent pooling layers reduce dimensionality while preserving salient features, and fully connected layers integrate this learned representation for downstream classification. CNNs have demonstrated strong performance across diverse genomics tasks, including TFBS classification, owing to their ability to robustly capture short-range dependencies and positional signals. However, their fixed receptive field constrains their ability to capture long-range interactions between distal regulatory elements, which are often critical for interpreting the regulatory mechanisms mediated by TFs [18]. Model training is generally performed using binary cross-entropy loss, with parameter optimization conducted via stochastic gradient descent or adaptive algorithms like Adam [44]. Despite their architectural simplicity, CNNs continue to serve as a foundational framework in genomic deep learning.

#### HyenaDNA

HyenaDNA [32] extends convolutional modeling to long-range genomic sequence processing using the Hyena operator [45]. In contrast to transformer architectures, which rely on self-attention, Hyena replaces it with a combination of structured long convolutions and data-controlled gating, enabling efficient modeling of both local and distal dependencies. A central innovation of HyenaDNA lies in its ability to dynamically generate convolutional filters as a function of sequence position. This allows the model to capture complex, position-specific interactions across extensive genomic contexts, achieving expressivity comparable to transformers but at significantly reduced computational cost. Importantly, HyenaDNA’s architecture scales sub-quadratically with input length, making it practical for processing sequences that span hundreds of thousands of base pairs. These characteristics make HyenaDNA particularly well-suited for large-scale genomic modeling tasks where long-range dependencies play a critical role, such as TF-DNA binding interactions.

#### Transformer-based Models

Transformer-based models have recently established themselves as the leading paradigm in DL for genomic sequence modeling [46]. Unlike CNNs, which are constrained by fixed receptive fields, transformers leverage self-attention mechanisms that allow each position in the input sequence to account for all the others. This enables the modeling of both short- and long-range dependencies, capturing information from large genomic contexts. Their ability to incorporate global context, makes transformers particularly well-suited for tasks where long-range interactions play a central role. By pretraining on large unlabeled genomic datasets, transformers learn rich representations of DNA sequences, which can then be fine-tuned for a variety of downstream applications, including TFBS prediction. In this study, we selected two representative models for benchmarking WaveDNA performance, DNABERT-2 and Nucleotide Transformer (NT), both of which have demonstrated competitive performance in TFBS prediction tasks.

#### DNABERT

DNABERT [25, 26] adapts the BERT architecture [47] to genomic sequences by employing the masked language modeling (MLM) objective. Input DNA sequences are tokenized into k-mers, embedded into high-dimensional vectors, and processed through stacked transformer encoder layers. These layers learn contextual representations by modeling dependencies across all positions in the sequence, allowing the model to capture both local and global regulatory patterns. DNABERT-2 introduces architectural improvements, that enhance the performance of the original model. Notably, it incorporates Attention with Linear Biases (ALiBi) [48], which improves generalization to sequences longer than those seen during training. Additionally, FlashAttention [29] is employed to reduce memory overhead and accelerate computation. DNABERT-2 replaces the fixed k-mer vocabulary with a variable-length vocabulary constructed via Byte-Pair Encoding (BPE), implemented through the SentencePiece tokenizer [49]. Together, these optimizations enhance both representational capacity and computational efficiency of the model, enabling DNABERT-2 to analyze longer genomic regions while achieving state-of-the-art performances across multiple genomic tasks, including TFBS prediction.

#### Nucleotide Transformer

Nucleotide Transformer (NT) [27] stands as one of the largest and most comprehensive transformer-based models for genomic sequence analysis. It employs an encoder-only architecture trained with the MLM objective. A key strength of NT lies in its large-scale pretraining across diverse genomic corpora, including human and multi-species genomic datasets. This extensive pretraining allows NT to capture both evolutionarily conserved elements and species-specific regulatory signatures, including TF-DNA binding patterns. NT achieves robust generalization and strong performances on multiple genomic tasks, including TFBS classification. Its robustness and broad applicability position NT as a foundational model in the field of genomic DL, serving as a versatile backbone for fine-tuning on specialized genomic prediction tasks.

## Results

We conducted a comprehensive evaluation of WaveDNA along three key aspects: predictive performance, computational efficiency, and biological interpretability. First, we benchmarked WaveDNA performance in TFBS prediction using 13 representative ChIP-seq datasets from ENCODE, employing the F1-score and Matthews Correlation Coefficient (MCC) as evaluation metrics [50]. Second, we examined the model’s computational scalability by measuring runtime and memory consumption, and compared its efficiency to that of HyenaDNA, DNABERT-2, and NT. Third, we investigated the biological interpretability of WaveDNA predictions through feature attribution analyses, comparing the resulting relevance maps with experimental ChIP-seq binding profiles. Our results show that WaveDNA achieves competitive or superior performance compared to transformer-based approaches, despite using ∼5 times fewer parameters and without relying on large-scale pretraining. Furthermore, WaveDNA demonstrates higher scalability and improved interpretability than transformer-based models. The interpretability analysis reveals that the features learned by WaveDNA align with experimentally validated binding signals, underscoring the model’s ability to capture biologically meaningful patterns driving TF-DNA binding

### Model fine-tuning for WaveDNA, HyenaDNA, DNABERT-2, and Nucleotide Transformer

For WaveDNA, fine-tuning was performed over three epochs using the AdamW optimizer [51] with a learning rate of 3 × 10™5 and a weight decay of 0.01. The number of training epochs (three) was empirically determined based on convergence of the training loss across all datasets (**Supplementary File Section 4**). Model optimization was guided by the cross-entropy loss, and training was conducted with a batch size of 4. WaveDNA was initialized with ResNet50 weights pretrained on ImageNet [52], provided by the torchvision library [53]. No additional pretraining or data augmentation procedures were applied. To ensure a fair and consistent comparison across architectures, DNABERT-2 and NT were fine-tuned using the same optimization scheme adopted for WaveDNA. The number of training epochs (three) was kept consistent across models, based on their convergence trends (**Supplementary Figure 3**). DNABERT-2 was fine-tuned with a batch size of 32, following the authors’ recommendations (https://github.com/MAGICS-LAB/DNABERT_2). No architectural modifications were introduced, and the original fine-tuning scripts were employed. In contrast, NT was trained using a batch size of 4, similar to WaveDNA, due to its larger memory footprint. The NT model family comprises architectures of varying sizes, ranging from hundreds of millions to billions of parameters, pretrained on genomic data from human and other species. We used the NT-500M variant, which encompasses approximately 500 million parameters and was pretrained on the human genome, representing a trade-off between model size and computational feasibility. The pretrained checkpoint was retrieved from the Hugging Face model hub (https://huggingface.co/Insta DeepAI/nucleotide-transformer-500m-human-rf). Unlike WaveDNA and DNABERT-2, which are explicitly designed for sequence classification tasks, NT follows a BERT-like architecture [47] that produces token-level embeddings. To adapt NT for TFBS prediction, we followed the protocol recommended by the authors: mean pooling was applied across token embeddings, followed by a fully connected multilayer perceptron classification head. This head consisted of two output neurons corresponding to TF-bound and unbound sequences, and was trained jointly with the transformer parameters. Finally, HyenaDNA was fine-tuned using the pretrained weights available at https://huggingface.co/LongSafari/hyenadna-medium-160k-seqlen-hf. All models, WaveDNA, HyenaDNA, DNABERT-2, and NT, were trained and evaluated on identical training and test dataset splits to ensure methodological consistency. All experiments were executed on a single NVIDIA GeForce RTX 4090 GPU (24 GB memory).

### WaveDNA Achieves Transformer-Level Performance in Genomic Sequence Classification Despite Smaller Size and No Pretraining on Genomic Sequences

We systematically evaluated the predictive performance of WaveDNA against HyenaDNA, DNABERT-2, and NT using F1-score and MCC across 13 TF datasets. The benchmarked models encompassed approximately 25 million parameters for WaveDNA, 117 million for DNABERT-2, 500 million for NT, and 6.5 million for HyenaDNA. Across chromosome-wise cross-validation experiments, WaveDNA consistently exhibited strong and class-balanced performance, achieving macro-averaged scores of F1 = 0.934 and MCC = 0.874. These values closely approached those of DNABERT-2 (F1 = 0.962, MCC = 0.924), while clearly surpassing NT (F1 = 0.879, MCC = 0.751) and markedly outperforming the less stable HyenaDNA baseline (F1 = 0.701, MCC = 0.485) (**Figure 2A** and **2B**). On average, WaveDNA trailed DNABERT-2 by ∼0.03 in F1 and ∼0.05 in MCC, yet exceeded NT by ∼0.05 F1 and ∼0.12 MCC, demonstrating near-parity with DNABERT-2 and a substantial advantage over NT. At individual TF level, WaveDNA outperformed NT across all 13 datasets and both metrics, with particularly pronounced gains for challenging TFs, such as JUND and GATA1. These represent the smallest datasets in the benchmark, containing the fewest labeled sequences and thus posing the most data-sparse prediction settings. Despite this, WaveDNA maintained high accuracy and stability, underscoring its robustness under limited data conditions. Although DNABERT-2 achieved marginally higher scores on certain datasets, WaveDNA matched or slightly surpassed it on others (e.g., JUND, F1: 0.962 vs. 0.956), and remained within a few thousandths on most remaining TFs (e.g., LCORL, RFX5). In contrast, HyenaDNA exhibited pronounced variability, performing competitively on a few TFs (e.g. ZBED1, TCF7L2) but showing marked lower performance on others. These results align with recent large-scale benchmarking studies reporting that transformer-based architectures, such as DNABERT-2 and Nucleotide Transformer consistently outperform HyenaDNA in genomic classification tasks [54, 55]. To statistically validate these findings, we conducted two-sample Kolmogorov–Smirnov tests to the distributions of F1 and MCC scores (**Figure 2C** and **2D**). Differences between WaveDNA and NT were statistically significant (p < 0.05) for both metrics, confirming the superior performance of WaveDNA. In contrast, no significant difference was observed between WaveDNA and DNABERT-2 (p > 0.1 for both F1 and MCC), indicating statistically comparable performance. We also conducted an ablation study to evaluate the impact of pretraining on WaveDNA by comparing models trained from scratch with those initialized using ImageNet-pretrained weights (**Supplementary File Section 5**). Pretrained initialization consistently enhanced performance across all metrics yielding higher performance (**Supplementary Table 2**), suggesting that pretraining provides a strong inductive bias that improves feature learning and convergence, particularly in data-limited settings. Collectively, these findings demonstrate that WaveDNA achieves performance comparable with DNABERT-2 while surpassing NT and HyenaDNA, despite operating without genomic pretraining and using approximately five times fewer parameters.

**Fig. 2:**
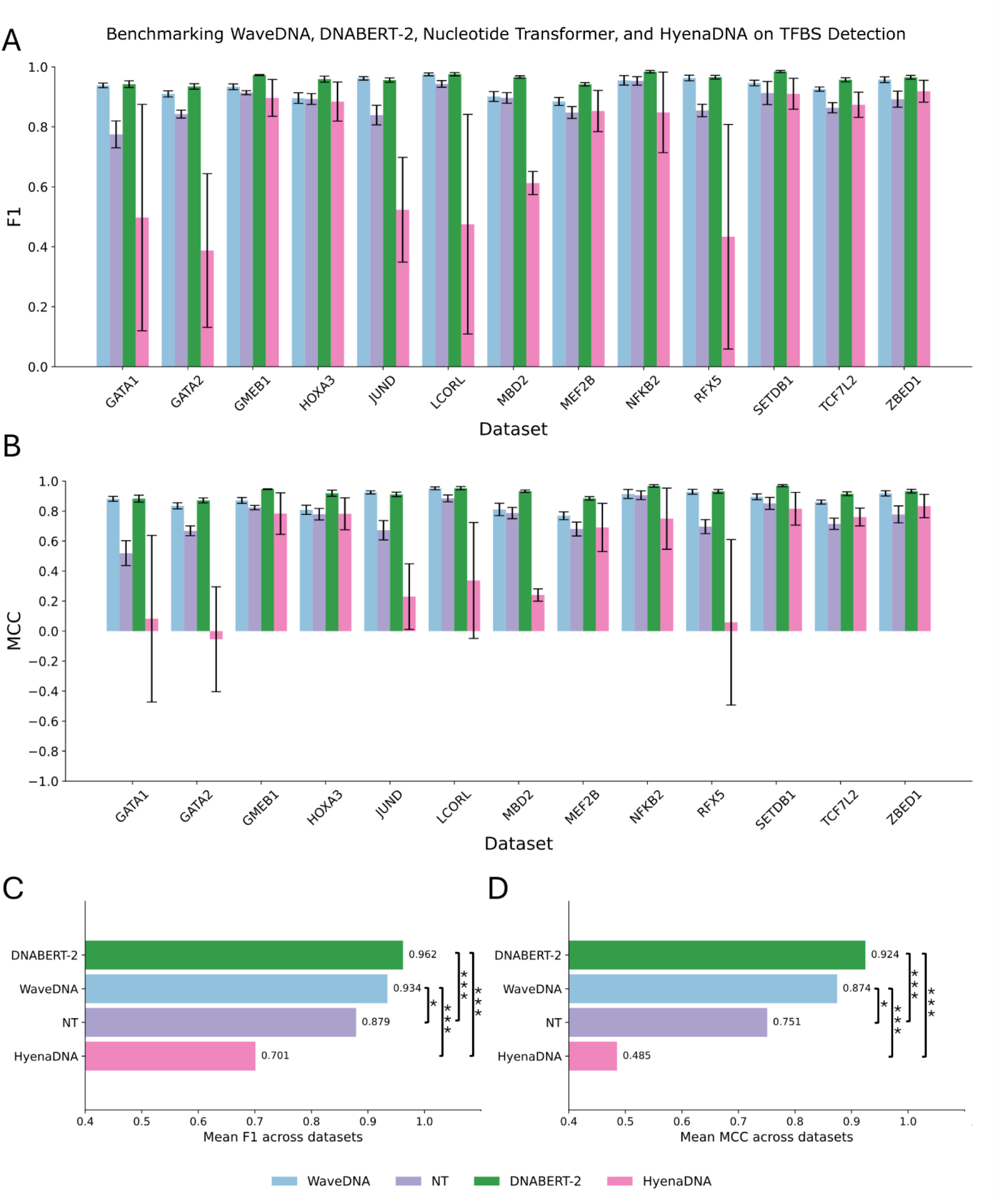
Performance comparison of WaveDNA, Nucleotide Transformer, DNABERT-2 and HyenaDNA. (A) F1 scores across 13 transcription factor datasets for WaveDNA (25M parameters), Nucleotide Transformer (500M), DNABERT-2 (117M), and HyenaDNA (6.5M). For each dataset and model, bars represent the mean F1 score across held-out chromosomes (1, 2, 3, 6, and 17), with error bars indicating the standard deviation. (B) Matthews Correlation Coefficient (MCC) values for the same datasets and models, reported as mean ± standard deviation across held-out chromosomes. (C–D) Model-level statistical comparison based on two-sample Kolmogorov–Smirnov tests applied to the distributions of (C) F1 and (D) MCC values across datasets. Asterisks denote significance levels (\**p <* 0.05, \*\**p <* 0.01, \*\*\**p <* 0.001).

### WaveDNA Exhibits Superior Scalability in Runtime and Memory Efficiency Compared to DNABERT-2 and Nucleotide Transformer

To evaluate models’ scalability and computational efficiency, we quantified both inference time and GPU memory utilization under controlled conditions (**Figure 3A** and **3B**). For runtime evaluation, we measured the average forward-pass duration per sequence across four input lengths (500, 1000, 2000, and 5000 base pairs) **(Figure 3A**). DNABERT-2 and NT displayed a superlinear increase in runtime with sequence length, consistent with the intrinsic quadratic complexity of their self-attention mechanisms. As a result, NT not only exhibited slower inference for longer inputs but also exceeded GPU memory limits when processing sequences longer than 2000 bp. In contrast, WaveDNA maintained an almost constant runtime across all tested input lengths, reflecting its architecture’s ability to project variable-length DNA sequences into fixed-size (224 × 224) image representations for ResNet50 processing. This design ensures consistent computational scalability even for long genomic sequences. Remarkably, despite this compression of sequence information into a fixed input size, WaveDNA preserved high predictive performance across different sequence lengths, underscoring the expressive capacity and robustness of the wavelet transform in extracting features from DNA sequences. While HyenaDNA achieved the lowest inference time overall, owing to its lightweight convolutional operators for long-range dependency modeling, this computational advantage comes at the expense of predictive accuracy (**Figure 2A** and **2B**). This trade-off highlights a key limitation of HyenaDNA, where efficiency gains coincide with diminished representational expressiveness. To further investigate scalability under varying computational loads, we profiled the GPU memory footprint during a single forward pass across batch sizes ranging from 4 to 128 on 1000 bp tokens sequences (**Figure 3B**). WaveDNA exhibited the smallest memory footprint across all tested configurations. At a batch size of 4, it required less than 1 GB of GPU memory, and even at batch size 128, its memory usage remained below 23 GB. DNABERT-2 and NT required substantially more memory due to their large parameter counts (117M and 500M, respectively) and the activation overhead typical of transformers. DNABERT-2 reached ∼18.5 GB of memory at batch size 16 and failed beyond that point, whereas NT already exceeded 13 GB at batch size 4 and could not scale further. Interestingly, despite having comparatively fewer parameters, HyenaDNA exhibited rapid memory growth as batch size increased, primarily due to its character-level tokenization strategy, which expands the dimensionality of intermediate activations. This effect led to out-of-memory conditions at batch size 128.

**Fig. 3:**
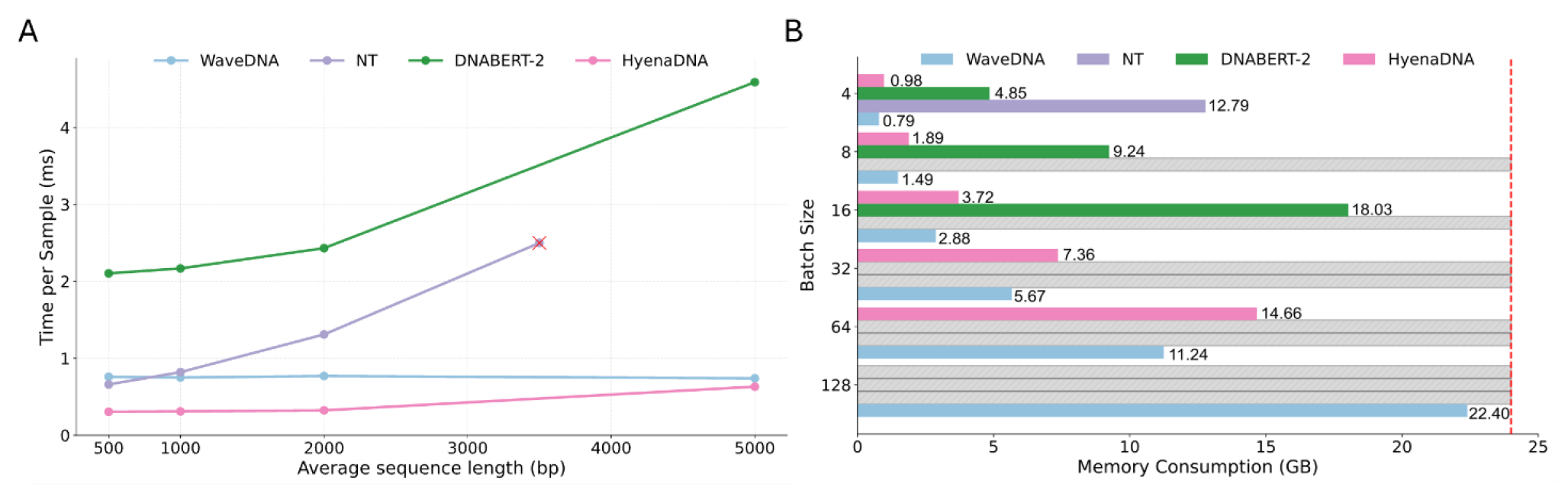
Scalability and computational efficiency of WaveDNA, Nucleotide Transformer, DNABERT-2 and HyenaDNA. (A) Runtime scalability of all models, measured as the average inference time per sequence across four input lengths (500, 1000, 2000, and 5000 bp). Red crosses indicate cases where models exceeded GPU memory capacity. (B) Memory efficiency of WaveDNA (25M parameters), Nucleotide Transformer (500M), DNABERT-2 (117M), and HyenaDNA (6.5M) quantified as GPU memory usage during a single forward pass across increasing batch sizes (4–128) with 1000 bp input sequences. The red dashed line marks the 24 GB GPU memory threshold, while gray bars highlight configurations that exceeded this limit

### WaveDNA Interpretability Through Convolutional Feature Attribution Correlates with ChIP-seq Evidence

To assess the interpretability of WaveDNA predictions and its ability to yield biologically meaningful insights, we employed Gradient-weighted Class Activation Mapping (Grad-CAM) (**Section 2.3**). This technique enables visualization of model attention by highlighting which regions of an input sequence contribute most to a specific classification decision. We focused our interpretability analysis on positive-class samples, i.e., bona fide TFBSs, since correlation with experimental ChIP-seq signals is meaningful only for true binding sites. For each test sample, Grad-CAM activations were computed using the convolutional layer within WaveDNA exhibiting the highest activation variance across the dataset. This method captures the most informative and diverse features across samples, thereby maximizing the interpretive value of the resulting activation maps. WaveDNA processes input DNA sequences through a wavelet transforms, yielding a two-dimensional representation where the x-axis encodes positional (time-domain) information along the sequence, while the y-axis corresponds to frequency components. Applying Grad-CAM to these feature maps produced 2D class activation maps, which were then reduced to 1D activation profiles by summing along the frequency axis, effectively condensing each map into a position-wise importance signal across the input sequence (**Figure 4**). To validate the biological relevance of activation signals, we retrieved bigWig tracks corresponding to the ChIP-seq experiment analyzed in this study from the ENCODE database. These tracks provide continuous quantitative measures of TF binding activity along genomic coordinates, with higher signal intensities (values) indicating stronger experimental binding evidence. Each bigwig file contains normalized reads density profiles for the corresponding TF, reflecting the experimentally measured binding strength across the genomic regions of interest. For each test sequence, we quantified the correlation between the Grad-CAM-derived activation map and the experimentally observed TF binding signal by computing the Pearson correlation coefficient between the Grad-CAM trace and the associated bigWig signal profile. This process was repeated across all test samples and datasets, yielding distributions of correlation values per TF. To evaluate whether these correlations were statistically significant, we performed one-sample t-tests against a null hypothesis of zero mean correlation. The results revealed statistically significant positive correlations (maximum p < 0.05) in all datasets), demonstrating that WaveDNA’s activation patterns align with experimentally measured TF binding signals (Figure 4). These findings suggest that WaveDNA’s predictions are grounded in biologically relevant sequence features rather than spurious dataset-specific correlations. In particular, the model consistently focused on regions coinciding with empirically validated binding signals. In contrast, transformers remain difficult to interpret. Their multi-head attention mechanisms, while powerful, do not always correlate with model importance or decision causality [56, 57, 58]. Moreover, the multilayer perceptron components of these models often suffer from feature superposition, where individual neurons encode multiple entangled and polysemantic features [59]. This property complicates direct interpretation and typically requires the training of auxiliary models to extract interpretable representation [60, 61]. By contrast, combining convolutional layers and Grad-CAM in WaveDNA, provides a spatially grounded attribution of model predictions to specific input regions. This provides a practical and interpretable framework for identifying functionally relevant sequence motifs or regulatory elements directly from model output.

**Fig. 4:**
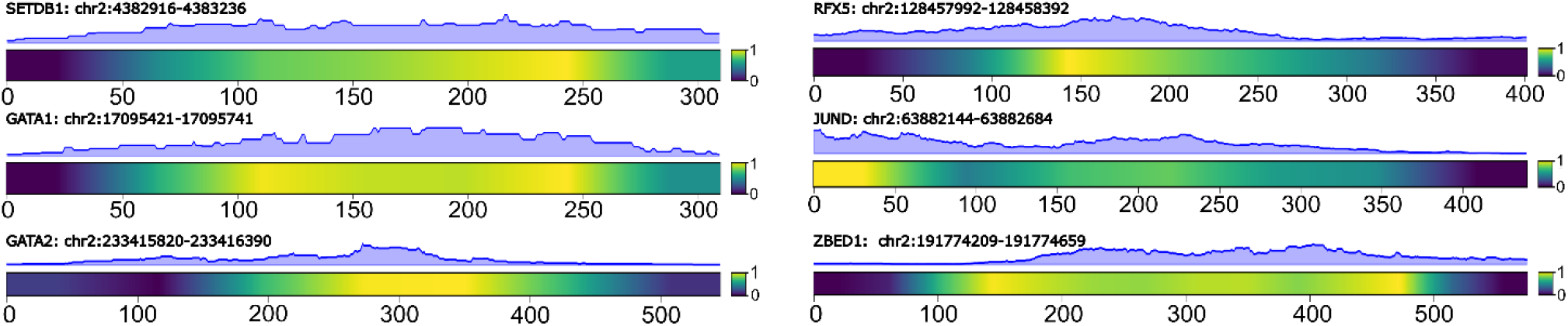
Interpretability of WaveDNA using Grad-CAM on test sequences from different transcription factors. The top panel shows the experimental ChIP-seq signal along a test DNA sequence, reflecting with transcription factor’s binding affinity. The bottom panel displays the Grad-CAM activation map computed by WaveDNA, obtained by summing activation values across image columns (frequency axis in the wavelet space) to produce a time-domain signal indicating each sequence position’s contribution to the final classification. Yellow regions correspond to positions where the model strongly focuses when predicting the positive class, while blue regions indicate low or negligible attention. High-activation regions align with ChIP-seq peaks, suggesting that WaveDNA effectively attends to biologically relevant sequence elements when making predictions.

## Conclusions

In this manuscript, we introduced WaveDNA, a novel framework based on wavelet transform for TFBS prediction. Through comprehensive evaluation on 13 ENCODE datasets, covering diverse DNA-binding domains, motif lengths, information content and sequence variability, we demonstrated that WaveDNA achieves performance comparable to transformer architectures, such as DNABERT-2, despite requiring significantly fewer parameters and no pretraining on genomic sequences. Furthermore, WaveDNA exhibited markedly improved scalability, maintaining constant runtime and lower memory consumption even at increasing input lengths, addressing a key limitation of transformer models. Critically, the convolutional architecture of WaveDNA enables spatially grounded attribution via Grad-CAM, producing feature maps that significantly correlate with experimental ChIP-seq signals. While this study focuses on TFBS prediction, the demonstrated effectiveness of WaveDNA in this task suggests that its architecture can serve as a viable and efficient alternative to more complex deep learning models for a broader range of genomic sequence classification problems. The framework could be extended to applications such as enhancer detection or splice site prediction, showcasing its versatility. Furthermore, representing DNA sequences as structured images opens new opportunities for generative modeling. Future work may explore the integration of diffusion-based architectures to generate realistic and biologically coherent DNA sequences, thereby bridging predictive modeling and generative genomics.

## Acknowledgments

The authors thank Prof. Luca Pinello, Prof. Erik Garrison and the members of InfOmics lab at University of Verona for their valuable suggestions and insightful discussion.

## Data Availability

WaveDNA code and detailed documentation to reproduce the results reported in this paper are available on GitHub (https://github.com/lorenzoruggerii/WaveDNA).

## Notes

### Competing Interest Statement

The authors have declared no competing interest.

### Summary of Updates

Results: Scalability Section Updated. Acknowledgments updated.

## References

1. Samuel A. Lambert, Arttu Jolma, Laura F. Campitelli, Pratyush K. Das, Yimeng Yin, Mihai Albu, Xiaoting Chen, Jussi Taipale, Timothy R. Hughes, and Matthew T. Weirauch. The human transcription factors. Cell, 172(4):650–665, 2018.

2. Alexander J. Stewart, Sridhar Hannenhalli, and Joshua B. Plotkin. Why transcription factor binding sites are ten nucleotides long. Genetics, 192(3): 973–985, 2012.

3. Matthew T. Maurano, Eric Haugen, Richard Sandstrom, Jeff Vierstra, Anthony Shafer, Rajinder Kaul, and John A. Stamatoyannopoulos. Large-scale identification of sequence variants influencing human transcription factor occupancy in vivo. Nature Genetics, 47(12): 1393–1401, 2015.

4. Manuel Tognon, Rosalba Giugno, and Luca Pinello. A survey on algorithms to characterize transcription factor binding sites. Briefings in Bioinformatics, 24(3):bbad156, 2023.

5. Michael F. Berger, Anthony A. Philippakis, Aaron M. Qureshi, Fangxue S. He, Preston W. Estep III, and Martha L. Bulyk. Compact, universal DNA microarrays to comprehensively determine transcription-factor binding site specificities. Nature Biotechnology, 24(11): 1429–1435, 2006.

6. Arttu Jolma, Teemu Kivioja, Jarkko Toivonen, Lu Cheng, Gonghong Wei, Martin Enge, Mikko Taipale, Juan M. Vaquerizas, Jian Yan, Mikko J. Sillanpää, et al. Multiplexed massively parallel SELEX for characterization of human transcription factor binding specificities. Genome Research, 20(6): 861–873, 2010.

7. David S. Johnson, Ali Mortazavi, Richard M. Myers, and Barbara Wold. Genome-wide mapping of in vivo protein-DNA interactions. Science, 316(5830): 1497–1502, 2007.

8. Sam John, Peter J. Sabo, Robert E. Thurman, Myong-Hee Sung, Simon C. Biddie, Thomas A. Johnson, Gordon L. Hager, and John A. Stamatoyannopoulos. Chromatin accessibility pre-determines glucocorticoid receptor binding patterns. Nature Genetics, 43(3): 264–268, 2011.

9. Ho Sung Rhee and B. Franklin Pugh. Comprehensive genome-wide protein-DNA interactions detected at single-nucleotide resolution. Cell, 147(6): 1408–1419, 2011.

10. Gary D. Stormo. DNA binding sites: representation and discovery. Bioinformatics, 16(1): 16–23, 2000.

11. Valentina Boeva. Analysis of genomic sequence motifs for deciphering transcription factor binding and transcriptional regulation in eukaryotic cells. Frontiers in Genetics, 7:24, 2016.

12. Manuel Tognon, Alisa Kumbara, Andrea Betti, Lorenzo Ruggeri, and Rosalba Giugno. Benchmarking transcription factor binding site prediction models: a comparative analysis on synthetic and biological data. Briefings in Bioinformatics, 26(4):bbaf363, 2025.

13. Ying He, Zhen Shen, Qinhu Zhang, Siguo Wang, and De-Shuang Huang. A survey on deep learning in DNA/RNA motif mining. Briefings in Bioinformatics, 22(4):bbaa229, 2021.

14. Haoyang Zeng, Matthew D. Edwards, Ge Liu, and David K. Gifford. Convolutional neural network architectures for predicting DNA–protein binding. Bioinformatics, 32(12):i121–i127, 2016.

15. Babak Alipanahi, Andrew Delong, Matthew T. Weirauch, and Brendan J. Frey. Predicting the sequence specificities of DNA- and RNA-binding proteins by deep learning. Nature Biotechnology, 33(8): 831–838, 2015.

16. David R. Kelley, Jasper Snoek, and John L. Rinn. Basset: learning the regulatory code of the accessible genome with deep convolutional neural networks. Genome Research, 26(7): 990–999, 2016.

17. Jian Zhou and Olga G. Troyanskaya. Predicting effects of noncoding variants with deep learning–based sequence model. Nature Methods, 12(10): 931–934, 2015.

18. Tianwei Yue, Yuanxin Wang, Longxiang Zhang, Chunming Gu, Haoru Xue, Wenping Wang, Qi Lyu, and Yujie Dun. Deep learning for genomics: A concise overview. arXiv preprint arXiv:1802.00810, 2018.

19. Sachi Inukai, Kian Hong Kock, and Martha L. Bulyk. Transcription factor–DNA binding: beyond binding site motifs. Current Opinion in Genetics & Development, 43: 110–119, 2017.

20. Gökcen Eraslan, Žiga Avsec, Julien Gagneur, and Fabian J. Theis. Deep learning: new computational modelling techniques for genomics. Nature Reviews Genetics, 20(7): 389–403, 2019.

21. Ashish Vaswani, Noam Shazeer, Niki Parmar, Jakob Uszkoreit, Llion Jones, Aidan N. Gomez, Łukasz Kaiser, and Illia Polosukhin. Attention is all you need. Advances in Neural Information Processing Systems, 30, 2017.

22. Žiga Avsec, Vikram Agarwal, Daniel Visentin, Joseph R. Ledsam, Agnieszka Grabska-Barwinska, Kyle R. Taylor, Yannis Assael, John Jumper, Pushmeet Kohli, and David R. Kelley. Effective gene expression prediction from sequence by integrating long-range interactions. Nature Methods, 18(10): 1196–1203, 2021.

23. Xi Fu, Shentong Mo, Alejandro Buendia, Anouchka P. Laurent, Anqi Shao, Maria del Mar Alvarez-Torres, Tianji Yu, Jimin Tan, Jiayu Su, Romella Sagatelian, et al. A foundation model of transcription across human cell types. Nature, 637(8047): 965–973, 2025.

24. David R. Kelley. Cross-species regulatory sequence activity prediction. PLoS Computational Biology, 16(7):e1008050, 2020.

25. Yanrong Ji, Zhihan Zhou, Han Liu, and Ramana V. Davuluri. DNABERT: pre-trained bidirectional encoder representations from transformers model for DNA-language in genome. Bioinformatics, 37(15): 2112–2120, 2021.

26. Zhihan Zhou, Yanrong Ji, Weijian Li, Pratik Dutta, Ramana Davuluri, and Han Liu. DNABERT-2: Efficient foundation model and benchmark for multi-species genome. arXiv preprint arXiv:2306.15006, 2023.

27. Hugo Dalla-Torre, Liam Gonzalez, Javier Mendoza-Revilla, Nicolas Lopez Carranza, Adam Henryk Grzywaczewski, Francesco Oteri, Christian Dallago, Evan Trop, Bernardo P. de Almeida, Hassan Sirelkhatim, et al. Nucleotide Transformer: building and evaluating robust foundation models for human genomics. Nature Methods, 22(2): 287–297, 2025.

28. Alexey Dosovitskiy, Lucas Beyer, Alexander Kolesnikov, Dirk Weissenborn, Xiaohua Zhai, Thomas Unterthiner, Mostafa Dehghani, Matthias Minderer, Georg Heigold, Sylvain Gelly, et al. An image is worth 16×16 words: Transformers for image recognition at scale. arXiv preprint arXiv:2010.11929, 2020.

29. Tri Dao, Dan Fu, Stefano Ermon, Atri Rudra, and Christopher Ré. FlashAttention: Fast and memory-efficient exact attention with IO-awareness. Advances in Neural Information Processing Systems, 35: 16344–16359, 2022.

30. Yi Tay, Mostafa Dehghani, Samira Abnar, Yikang Shen, Dara Bahri, Philip Pham, Jinfeng Rao, Liu Yang, Sebastian Ruder, and Donald Metzler. Long Range Arena: A benchmark for efficient transformers. arXiv preprint arXiv:2011.04006, 2020.

31. Jim Clauwaert, Gerben Menschaert, and Willem Waegeman. Explainability in transformer models for functional genomics. Briefings in Bioinformatics, 22(5):bbab060, 2021.

32. Eric Nguyen, Michael Poli, Marjan Faizi, Armin Thomas, Michael Wornow, Callum Birch-Sykes, Stefano Massaroli, Aman Patel, Clayton Rabideau, Yoshua Bengio, et al. HyenaDNA: Long-range genomic sequence modeling at single nucleotide resolution. Advances in Neural Information Processing Systems, 36: 43177–43201, 2023.

33. Sasha Targ, Diogo Almeida, and Kevin Lyman. ResNet in ResNet: Generalizing residual architectures. arXiv preprint arXiv:1603.08029, 2016.

34. ENCODE Project Consortium et al. An integrated encyclopedia of DNA elements in the human genome. Nature, 489(7414): 57, 2012.

35. Ramprasaath R. Selvaraju, Michael Cogswell, Abhishek Das, Ramakrishna Vedantam, Devi Parikh, and Dhruv Batra. Grad-CAM: Visual explanations from deep networks via gradient-based localization. In Proceedings of the IEEE International Conference on Computer Vision, pages 618–626, 2017.

36. Olivier Rioul and Pierre Duhamel. Fast algorithms for discrete and continuous wavelet transforms. IEEE Transactions on Information Theory, 38(2): 569–586, 2002.

37. Christopher M. Leavey, M. Neil James, John Summerscales, and Robert Sutton. An introduction to wavelet transforms: a tutorial approach. Insight—Non-Destructive Testing and Condition Monitoring, 45(5): 344–353, 2003.

38. Kaiming He, Xiangyu Zhang, Shaoqing Ren, and Jian Sun. Deep residual learning for image recognition. In Proceedings of the IEEE Conference on Computer Vision and Pattern Recognition, pages 770–778, 2016.

39. Bolei Zhou, Aditya Khosla, Agata Lapedriza, Aude Oliva, and Antonio Torralba. Learning deep features for discriminative localization. In Proceedings of the IEEE Conference on Computer Vision and Pattern Recognition, pages 2921–2929, 2016.

40. Wouter Meuleman, Alexander Muratov, Eric Rynes, Jessica Halow, Kristen Lee, Daniel Bates, Morgan Diegel, Douglas Dunn, Fidencio Neri, Athanasios Teodosiadis, et al. Index and biological spectrum of human DNase I hypersensitive sites. Nature, 584(7820): 244–251, 2020.

41. Ning Sun and Hongyu Zhao. Reconstructing transcriptional regulatory networks through genomics data. Statistical Methods in Medical Research, 18(6): 595–617, 2009.

42. Jason Gertz, Daniel Savic, Katherine E. Varley, E. Christopher Partridge, Alexias Safi, Preti Jain, Gregory M. Cooper, Timothy E. Reddy, Gregory E. Crawford, and Richard M. Myers. Distinct properties of cell-type-specific and shared transcription factor binding sites. Molecular Cell, 52(1): 25–36, 2013.

43. Felix Teufel, Magnús Halldór Gíslason, José Juan Almagro Armenteros, Alexander Rosenberg Johansen, Ole Winther, and Henrik Nielsen. GraphPart: homology partitioning for biological sequence analysis. NAR Genomics and Bioinformatics, 5(4):lqad088, 2023.

44. Diederik P. Kingma and Jimmy Ba. Adam: A method for stochastic optimization. arXiv preprint arXiv:1412.6980, 2017.

45. Michael Poli, Stefano Massaroli, Eric Nguyen, Daniel Y. Fu, Tri Dao, Stephen Baccus, Yoshua Bengio, Stefano Ermon, and Christopher Ré. Hyena Hierarchy: Towards larger convolutional language models. In International Conference on Machine Learning, pages 28043–28078. PMLR, 2023.

46. Shuang Zhang, Rui Fan, Yuti Liu, Shuang Chen, Qiao Liu, and Wanwen Zeng. Applications of transformer-based language models in bioinformatics: a survey. Bioinformatics Advances, 3(1):vbad001, 2023.

47. Jacob Devlin, Ming-Wei Chang, Kenton Lee, and Kristina Toutanova. BERT: Pre-training of deep bidirectional transformers for language understanding. In Proceedings of the 2019 Conference of the North American Chapter of the Association for Computational Linguistics: Human Language Technologies, volume 1 (Long and Short Papers), pages 4171–4186, 2019.

48. Ofir Press, Noah A. Smith, and Mike Lewis. Train short, test long: Attention with linear biases enables input length extrapolation. arXiv preprint arXiv:2108.12409, 2021.

49. Taku Kudo and John Richardson. SentencePiece: A simple and language-independent subword tokenizer and detokenizer for neural text processing. arXiv preprint arXiv:1808.06226, 2018.

50. Davide Chicco and Giuseppe Jurman. The advantages of the Matthews correlation coefficient (MCC) over F1 score and accuracy in binary classification evaluation. BMC Genomics, 21(1): 6, 2020.

51. Ilya Loshchilov and Frank Hutter. Decoupled weight decay regularization. arXiv preprint arXiv:1711.05101, 2017.

52. Jia Deng, Wei Dong, Richard Socher, Li-Jia Li, Kai Li, and Li Fei-Fei. ImageNet: A large-scale hierarchical image database. In 2009 IEEE Conference on Computer Vision and Pattern Recognition, pages 248–255. IEEE, 2009.

53. Sébastien Marcel and Yann Rodriguez. Torchvision: The machine-vision package of Torch. In Proceedings of the 18th ACM International Conference on Multimedia, pages 1485–1488, 2010.

54. Haonan Feng, Lang Wu, Bingxin Zhao, Chad Huff, Jianjun Zhang, Jia Wu, Lifeng Lin, Peng Wei, and Chong Wu. Benchmarking DNA foundation models for genomic sequence classification. bioRxiv, 2024.

55. Zicheng Liu, Jiahui Li, Siyuan Li, Zelin Zang, Cheng Tan, Yufei Huang, Yajing Bai, and Stan Z. Li. GenBench: A benchmarking suite for systematic evaluation of genomic foundation models. arXiv preprint arXiv:2406.01627, 2024.

56. Sofia Serrano and Noah A. Smith. Is attention interpretable? arXiv preprint arXiv:1906.03731, 2019.

57. Bing Bai, Jian Liang, Guanhua Zhang, Hao Li, Kun Bai, and Fei Wang. Why attentions may not be interpretable? In Proceedings of the 27th ACM SIGKDD Conference on Knowledge Discovery & Data Mining, pages 25–34, 2021.

58. Sarah Wiegreffe and Yuval Pinter. Attention is not not explanation. arXiv preprint arXiv:1908.04626, 2019.

59. Nelson Elhage, Tristan Hume, Catherine Olsson, Nicholas Schiefer, Tom Henighan, Shauna Kravec, Zac Hatfield-Dodds, Robert Lasenby, Dawn Drain, Carol Chen, et al. Toy models of superposition. arXiv preprint arXiv:2209.10652, 2022.

60. Hoagy Cunningham, Aidan Ewart, Logan Riggs, Robert Huben, and Lee Sharkey. Sparse autoencoders find highly interpretable features in language models. arXiv preprint arXiv:2309.08600, 2023.

61. Jacob Dunefsky, Philippe Chlenski, and Neel Nanda. Transcoders find interpretable LLM feature circuits. Advances in Neural Information Processing Systems, 37: 24375–24410, 2024.

