## Supplementary material for "Improving DNA Modeling with WaveDNA: Enhancing Speed, Generalizability, and Interpretability through Wavelet Transformation": Supplemetary File

#### 1 Datasets Selection and Processing

In this study, we evaluated the models’ performance on ChIP-seq data for 13 distinct transcription factors (TFs) representing diverse DNA-binding domains [1], using optimal IDR-thresholded peaks obtained from the ENCODE database [2]. The ChIP-seq data were provided in the narrowPeak format, an ENCODE-specific BED format commonly used for TFs due to their typically narrow binding regions.

To extract the corresponding DNA sequences in FASTA format, we employed the *getfasta* command from BEDTools [3], which retrieves sequences defined by BED intervals coordinates and generates matching FASTA entries using the supplied reference genome.

For background data, we downloaded DNase-seq peaks in BED format from ENCODE [4]. To ensure non-overlapping background sequences, we used BEDTools’ *intersect* and *subtract* commands to remove any DNase-seq feature that overlapped with ChIP-seq peaks. This procedure generated a clean set of DNase-seq regions suitable for negative/background sequences.

To rigorously evaluate model generalization and minimize the risk of overfitting chromosome-specific sequence patterns, we implemented a chromosome-based cross-validation strategy across all TF datasets. For each TF, sequences mapped on chromosomes 1, 2, 3, 6, and 17 were sequentially held out as independent test sets while remaining chromosomes’ sequences were used for model training. These chromosomes were selected because they contained the largest numbers of positive and negative sequences in each dataset, providing representative and balanced test subsets (**Supplementary Figures 1** and **2**).

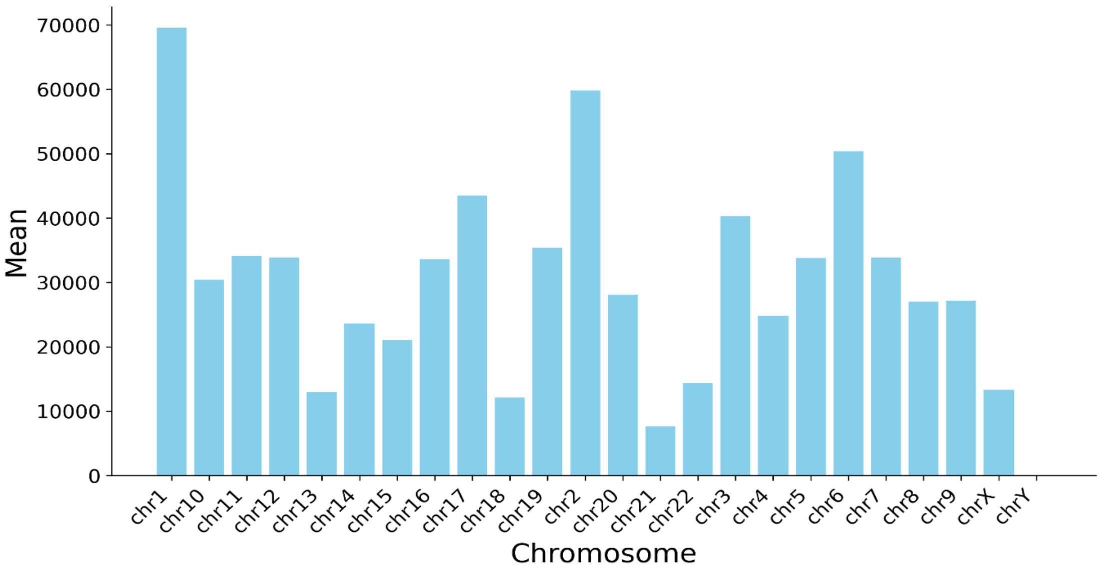


**Supplementary Figure 1:** Average number of sequences per chromosome across the 13 transcription factor (TF) datasets. The figure shows the mean number of sequences mapped to each chromosome for all the 13 ChIP-seq TF datasets analyzed. Chromosomes 1, 2, 3, 6, and 17 consistently rank among the top five in sequence abundance across nearly all TFs.

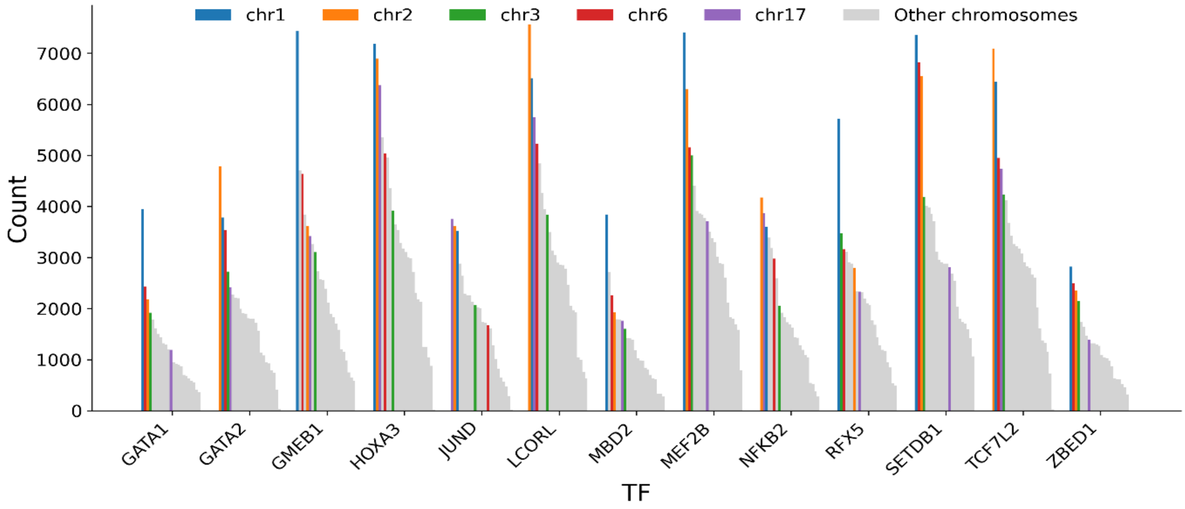


**Supplementary Figure 2:** Chromosomal distribution of sequences across transcription factor (TF) datasets. The figure illustrates the distribution of sequences per chromosome for each TF dataset used in model training and evaluation. Chromosomes 1, 2, 3, 6, and 17 (highlighted in the figure) consistently display high sequence abundance.

### 2 Assessment of Sequence Homology between Training and Test Datasets

To address the potential confounding effects of homologous sequences that could bias model training and artificially inflate performance metrics [5,6], we evaluated sequence similarity between training and test sets.

We systematically assessed sequence homology between the two datasets using pairwise global alignment based on the Needleman-Wunsch algorithm [7], as implemented in Biopython [8]. This method computes optimal alignments over entire sequence lengths, providing a robust measure of overall similarity. For each test sequence, the highest percentage of identity with any training sequence was recorded according to the best-scoring alignment, ensuring a rigorous assessment of potential data leakage.

We applied a conservative homology threshold of 80% sequence identity, considering any test sequence exceeding this threshold as potentially non-independent. This cutoff is widely adopted to denote substantial sequence similarity, which can result in model overfitting and artificially inflated predictive performance [5,6].

To quantify the extent and impact of sequence homology, we calculated the proportion of test sequences surpassing the 80% identity threshold relative to the training dataset. The analysis was further stratified into test-positive and test-negative subsets to determine whether homology-induced data leakage disproportionately affected model evaluation across distinct sequence classes.

Across all datasets, sequence homology remained consistently low, not even reaching 1% (**Supplementary Table 1**). These results show that no significant data leakage occurred between training and testing sets, supporting robustness and independence of data splits used for benchmarking models' performance.

### 3 Benchmarked Models

In this section, we provide a detailed description of the architectures of the benchmarked models employed in this study.

CNN-based architectures for transcription factor binding site (TFBS) prediction typically comprise convolutional, pooling, and fully connected layers, complemented by regularization mechanisms [9,10]. Input DNA sequences are one-hot encoded into matrices $X \in \{0,1\}^{4\times|s|}$, which are processed by learnable convolutional kernels $\{W^{(k)} \in \mathbb{R}^{4\times w}\}_{k=1}^{K}$ to generate activation maps $z^{(k)}$ that capture local motif-like patterns. Subsequent max-pooling reduces dimensionality and highlights salient features, which are then integrated by fully connected layers with dropout regularization. The final layer applies a sigmoid or softmax function to produce class probabilities, optimized using binary cross-entropy loss and stochastic or adaptive gradient-based methods such as Adam [11]. The HyenaDNA architecture [12] adopts a decoder-only sequence-to-sequence framework composed of alternating Hyena operator blocks and feed-forward layers. Each Hyena operator [13] replaces self-attention with a structured combination of long convolutions and data-controlled gating, constructing diagonal gating matrices $D_{x_1}, D_{x_2}$ and a Toeplitz convolutional matrix $T_h$ parameterized by a small neural network $\gamma_\theta$. This design enables efficient modeling of long-range dependencies with $O(L\log_2 L)$ complexity, allowing scalability to genomic sequences spanning hundreds of thousands of base pairs.

DNABERT [14,15], derived from the BERT architecture [16], tokenizes DNA into k-mers represented as embedding vectors forming input matrices $M \in \mathbb{R}^{T\times d}$. It consists of 12 transformer encoder layers, each with 12 attention heads and a hidden dimension of 768, where each attention head computes contextual dependencies via the softmax-weighted dot product of projected queries, keys, and values. DNABERT variants differ by k-mer size ($k \in \{3,4,5,6\}$), while DNABERT-2 introduces several architectural enhancements for scalability and efficiency, including Attention with Linear Biases (ALiBi) [17] for relative positional encoding, FlashAttention [18] for memory-efficient exact attention computation, and Low-Rank Adaptation (LoRA) [19] for parameter-efficient fine-tuning. Unlike DNABERT, which employs fixed-length k-mers, DNABERT-2 utilizes variable-length subword tokenization based on Byte-Pair Encoding (BPE) via the SentencePiece tokenizer [20], reducing the number of tokens per sequence and mitigating the quadratic complexity of attention.

The Nucleotide Transformer (NT) [21] represents a family of large-scale encoder-only transformer models pretrained on genomic sequences using the Masked Language Modeling (MLM) objective. Input sequences are tokenized into non-overlapping 6-mers and embedded into vectors with learned positional encodings to preserve sequence order. The model comprises stacked transformer encoder layers with multi-head self-attention and feed-forward networks employing GELU activations, residual connections, and layer normalization, enabling hierarchical representations that capture both local and global dependencies. The NT collection encompasses models of varying parameter scales trained on human-specific and multi-species genomic data, supporting a wide range of downstream genomics applications, including TFBS prediction.

### 4 Training Loss Analysis

To assess the learning dynamics of the benchmarked models (WaveDNA, HyenaDNA, DNABERT-2, and NT), we examined the evolution of their training loss across epochs. Training was conducted using different batch sizes (WaveDNA, HyenaDNA, and NT: 4; DNABERT-2: 32) to accommodate the varying computational requirements and architectural characteristics of each model. Across all architectures, the mean training loss exhibits a consistent downward trend during the initial epochs, followed by stabilization after approximately three epochs, corresponding to 100% of the training progression (**Supplementary Figure 3**). This convergence behavior indicates that all models effectively learn stable representations from genomic data and that the chosen optimization configurations support efficient and robust training dynamics across datasets.

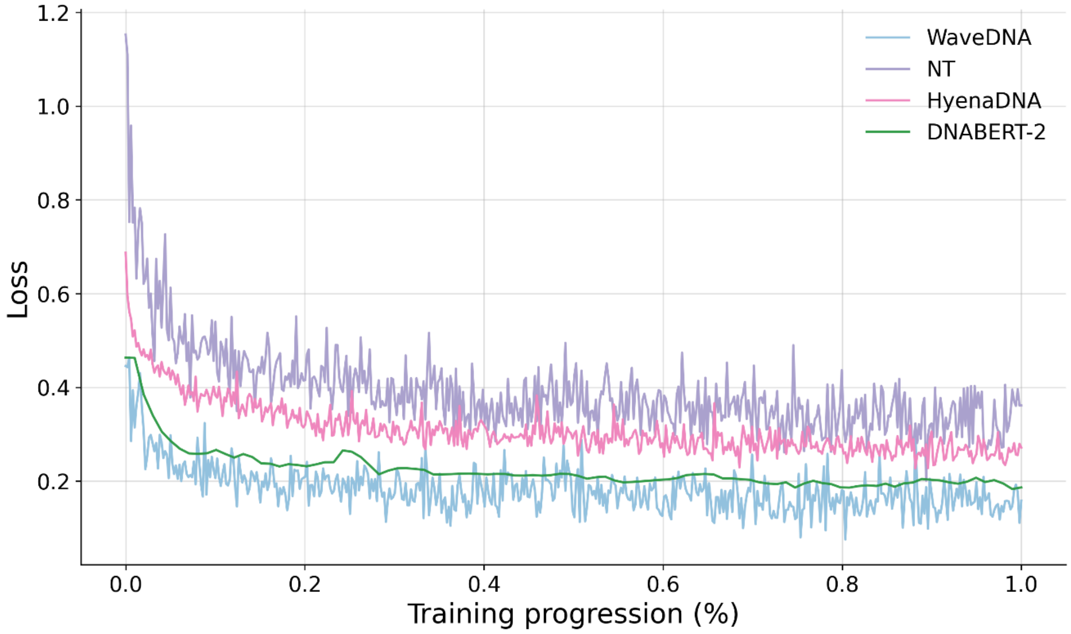


**Supplementary Figure 3:** Mean training loss across epochs for WaveDNA, HyenaDNA, DNABERT-2, and NT. Training was performed with different batch sizes (WaveDNA, HyenaDNA and NT: 4, DNABERT-2: 32). Across all models, the mean training loss stabilizes after three epochs (corresponding to 100% training progression), indicating stable and effective learning across datasets.

### 5 Ablation Studies

To evaluate the contribution of pretraining to WaveDNA's predictive performance, we performed an ablation study comparing two configurations: (i) models trained from scratch, and (ii) models initialized with weights pretrained on the ImageNet dataset [22].

The results indicate that pretraining on ImageNet substantially enhances performance across all evaluation metrics (**Supplementary Table 2**). Specifically, models initialized with pretrained weights achieved significantly higher F1 and Matthews Correlation Coefficient (MCC) scores [23] ($p$ < 0.05, KS test), reflecting both improved sensitivity and overall classification reliability. Pretraining yielded faster convergence and more stable training dynamics, suggesting that the low-level features captured by CNNs in natural images transfer effectively to the genomic domain when DNA sequences are represented as time-frequency images. This transfer learning setup provides a strong inductive bias, facilitating more efficient feature extraction and regularization, particularly in low-data regimes.

Notably, the effect was most pronounced for smaller datasets (e.g., GATA1), where pretrained initialization mitigated overfitting and enhanced generalization. Overall, these results highlight the benefit of incorporating image-based pretraining in WaveDNA, demonstrating that knowledge transfer from computer vision can meaningfully improve model robustness and accuracy in genomic sequence modeling tasks.

### References


1. Samuel A Lambert, Arttu Jolma, Laura F Campitelli, Pratyush K Das, Yimeng Yin, Mihai Albu, Xiaoting Chen, Jussi Taipale, Timothy R Hughes, and Matthew T Weirauch. The human transcription factors. *Cell*, 172(4):650—665, 2018.
2. ENCODE Project Consortium et al. An integrated encyclopedia of dna elements in the human genome. *Nature*, 489(7414):57, 2012.
3. Aaron R Quinlan, Ira M. Hall. BEDTools: a flexible suite of utilities for comparing genomic features. *Bioinformatics*, 26(6):841—842, 2010.
4. Wouter Meuleman, Alexander Muratov, Eric Rynes, Jessica Harlow, Kristen Lee, Daniel Bates, Morgan Diegel, Douglas Dunn, Fidencio Neri, Athanasios Teodosiadis, et al. Index and biological spectrum of human DNase I hypersensitive sites. *Nature* 584(7820):244—251, 2020.
5. Felix Teufel, Magnús Halldór Gíslason, José Juan Almagro Armenteros, Alexander Rosenberg Johansen, Ole Winther, and Henrik Nielsen. GraphPart: homology partitioning for biological sequence analysis. *NAR genomics and bioinformatics*, 5(4):lqad088, 2023.
6. Abdul Muntakim Rafi, Brett Kiyota, Nozomu Yachie, Carl de Boer. Detecting and avoiding homology-based data leakage in genome-trained sequence models. *bioRxiv*, 2025.
7. Richard Durbin, Sean R. Eddy, Anders Krogh, Graeme Mitchison. Biological sequence analysis: probabilistic models of proteins and nucleic acids. Cambridge university press 1998.
8. Peter J Cock, Tiago Antao, Jeffrey T Chang, Brad A Chapman, Cymon J Cox, Andrew Dalke, Iddo Friedberg, Thomas Hamelryck, Frank Kauff, Bartek Wilczynski, et al. Biopython: freely available Python tools for computational molecular biology and bioinformatics. *Bioinformatics*, 25(11):1422—1423, 2009.
9. Manuel Tognon, Rosalba Giugno, and Luca Pinello. A survey on algorithms to characterize transcription factor binding sites. *Briefings in Bioinformatics*, 24(3):bbad156, 2023.
10. Haoyang Zeng, Matthew D Edwards, Ge Liu, and David K Gifford. Convolutional neural network architectures for predicting dna–protein binding. *Bioinformatics*, 32(12):i121–i127, 2016.
11. Diederik P. Kingma and Jimmy Ba. Adam: A method for stochastic optimization. *arXiv preprint arXiv*:1412.6980, 2017.
12. Eric Nguyen, Michael Poli, Marjan Faizi, Armin Thomas, Michael Wornow, Callum Birch-Sykes, Stefano Massaroli, Aman Patel, Clayton Rabideau, Yoshua Bengio, et al. Hyenadna: Long-range genomic sequence modeling at single nucleotide resolution. *Advances in neural information processing systems*, 36:43177—43201, 2023.
13. Michael Poli, Stefano Massaroli, Eric Nguyen, Daniel Y Fu, Tri Dao, Stephen Baccus, Yoshua Bengio, Stefano Ermon, and Christopher Ré. Hyena hierarchy: Towards larger convolutional language models. In *International Conference on Machine Learning*, pages 28043–28078. PMLR, 2023
14. Yanrong Ji, Zhihan Zhou, Han Liu, and Ramana V Davuluri. Dnabert: pre-trained bidirectional encoder representations from transformers model for dna-language in genome. *Bioinformatics*, 37(15):2112—2120, 2021.
15. Zhihan Zhou, Yanrong Ji, Weijian Li, Pratik Dutta, Ramana Davuluri, and Han Liu. Dnabert-2: Efficient foundation model and benchmark for multi-species genome. *arXiv preprint arXiv*:2306.15006, 2023
16. Jacob Devlin, Ming-Wei Chang, Kenton Lee, and Kristina Toutanova. BERT: Pre-training of deep bidirectional transformers for language understanding. In *Proceedings of the 2019 Conference of the North American Chapter of the Association for Computational Linguistics: Human Langiage Technologies* 1:4171—4186, 2019.

17. Ofir Press, Noah A Smith, and Mike Lewis. Train short, test long: Attention with linear biases enables input length extrapolation. *arXiv preprint arXiv*: 2108.12409, 2021.
18. Tri Dao. Flashattention-2: Faster attention with better parallelism and work partitioning. *arXiv preprint arXiv*:2307.08691, 2023.
19. Edward J Hu, Yelong Shen, Philip Wallis, Zeyuan Allen-Zhu, Yuanzhi Li, Shean Wang, Lu Wang, and Weizhu Chen. Lora: Low-rank adaptation of large language models. *ICLR* 1(2):3, 2022.
20. Taku Kudo and John Richardson. SentencePiece: A simple and language independent subword tokenizer and detokenizer for neural text processing. *arXiv preprint arXiv*:1808.06226, 2018.
21. Hugo Dalla-Torre, Liam Gonzalez, Javier Mendoza-Revilla, Nicolas Lopez Carranza, Adam Henryk Grzywaczewski, Francesco Oteri, Christian Dallago, Evan Trop, Bernardo P de Almeida, Hassan Sirelkhatim, et al. Nucleotide transformer: building and evaluating robust foundation models for human genomics. *Nature Methods*, 22(2):287–297, 2025.
22. Jia Deng, Wei Dong, Richard Socher, Li-Jia Li, Kai Li, and Li Fei-Fei. Imagenet: A large-scale hierarchical image database. In 2009 IEEE conference on computer vision and pattern recognition, pages 248–255. Ieee, 2009.
23. Davide Chicco and Giuseppe Jurman. The advantages of the matthews correlation coefficient (mcc) over f1 score and accuracy in binary classification evaluation. BMC genomics, 21(1):6, 2020.

| Experiment | Target | Homology Percentage (%) | Positive Homology Percentage (%) | Negative Homology Percentage (%) |
|---|---|---|---|---|
| ENCFF365ETH | GMEB1 | 0,056721 | 0,113443 | 0 |
| ENCFF409YKQ | GATA2 | 0,272308 | 0,502723 | 0,041894 |
| ENCFF438ZEN | RFX5 | 0 | 0 | 0 |
| ENCFF606UCO | LCORL | 0,039714 | 0,052952 | 0,026476 |
| ENCFF657CTC | GATA1 | 0,091827 | 0,183655 | 0 |
| ENCFF658HLG | JUND | 0 | 0 | 0 |
| ENCFF687TJX | NFKB2 | 0,095877 | 0,191755 | 0 |
| ENCFF726SWQ | TCF7L2 | 0,45403 | 0,908059 | 0 |
| ENCFF753OCA | HOXA3 | 0,364857 | 0,671337 | 0,058377 |
| ENCFF753UAX | ZBED1 | 0 | 0 | 0 |
| ENCFF788YHU | MBD2 | 0 | 0 | 0 |
| ENCFF812YDP | SETDB1 | 0,753532 | 1,507064 | 0 |
| ENCFF884QQW | MEF2B | 0,016683 | 0 | 0,033367 |

**Supplementary Table 1.** Summary of sequence homology across the 13 Chip-seq experiments used for models' performance evaluation. For each experiment, the table reports the proportion of homologous sequences in the test set, including the overall percentage as well as separate values for positive and negative test subsets.

| TF Dataset | Ablation (MCC) | No Ablation (MCC) |
|---|---|---|
| GATA1 | 0.2467 ± 0.0623 | **0.8810 ± 0.0161** |
| GATA2 | 0.7139 ± 0.0708 | **0.8651 ± 0.0208** |
| GMEB1 | 0.6897 ± 0.0482 | **0.8709 ± 0.0190** |
| HOXA3 | 0.6711 ± 0.0482 | **0.8081 ± 0.0308** |
| JUND | 0.7118 ± 0.0284 | **0.9246 ± 0.0103** |
| LCORL | 0.7938 ± 0.0377 | **0.9516 ± 0.0094** |
| MBD2 | 0.7292 ± 0.0299 | **0.8109 ± 0.0412** |
| MEF2B | 0.6333 ± 0.0377 | **0.7691 ± 0.0259** |
| NFKB2 | 0.7561 ± 0.0596 | **0.9140 ± 0.0297** |
| RFX5 | 0.7491 ± 0.0799 | **0.9284 ± 0.0166** |
| SETDB1 | 0.7579 ± 0.0486 | **0.8955 ± 0.0193** |
| TCF7L2 | 0.7351 ± 0.0359 | **0.8598 ± 0.0134** |
| ZBED1 | 0.6518 ± 0.0623 | **0.9180 ± 0.0171** |

**Supplementary Table 2:** Comparison of WaveDNA performance with and without ImageNet pretraining. Reported are mean ± standard deviation of MCC scores across chromosomes for models trained from scratch (Ablation) and models initialized with ImageNet-pretrained weights (No Ablation).
